# Multilocus Sequence Typing Reveals A Unique Co-Dominant Population Structure of *Cryptococcus Neoformans* Var. *Grubii* in Vietnam

**DOI:** 10.1101/190785

**Authors:** Thanh Lam Tuan, Trieu Phan Hai, Sayaphet Rattanavong, Trinh Mai Nguyen, Anh Duong Van, Cherrelle Dacon, Thu Hoang Nha, Lan Phu Huong Nguyen, Chau Thi Hong Tran, Viengmon Davong, Chau Van Vinh Nguyen, Guy E. Thwaites, Maciej F. Boni, David Dance, Philip M. Ashton, Jeremy N. Day

## Abstract

Cryptococcosis is amongst the most important invasive fungal infections globally, with cryptococcal meningitis causing an estimated 180,000 deaths each year in HIV infected patients alone. Patients with other forms of immunosuppression are also at risk, and disease is increasingly recognized in apparently immunocompetent individuals. *Cryptococcus neoformans* var. *grubii* (serotype A, molecular type VNI) has a global distribution and is responsible for the majority of cases. Here, we used the consensus ISHAM Multilocus Sequence Typing (MLST) for *C. neoformans* to define the population structure of clinical isolates of *Cryptococcus neoformans* var. *grubii* from Vietnam (n=136) and Laos (n=81). We placed these isolates into the global context using published MLST data from 8 other countries (total N = 669). We observed a phylo-geographical relationship in which Laos was similar to its Southeast Asian neighbor Thailand in being dominated (83%) by Sequence Type (ST) 4 and its Single Locus Variant ST6. On the other hand, Vietnam was uniquely intermediate between Southeast Asia and East Asia having both ST4/ST6 (35%) and ST5 (48%) which causes the majority of cases in East Asia. Analysis of genetic distance (Fst) between different populations of *Cryptococcus neoformans* var. *grubii* supported the intermediate nature of the population from Vietnam. A strong association between ST5 and infection in apparently immunocompetent, HIV-uninfected patients was observed in Vietnam (OR: 7.97, [95%CI: 3.18-19.97], *p* < 0.0001). Our study emphasizes that Vietnam, with its intermediate *Cryptococcus neoformans* var. *grubii* population structure, provides the strongest epidemiological evidence of the relationship between ST5 and infection of HIV-uninfected patients. Human population genetic distances within the region
suggest these differences in CNVG population across Southeast Asia are driven by ecological factors rather than host factors.

**Author summary:** *Cryptococcus neoformans* is a yeast that causes meningitis in people, usually with damaged immune systems. There are >180,000 deaths in HIV-infected patients each year, most occurring where there are the highest HIV/AIDS disease burdens. Vietnam and Laos have contributed significantly to clinical trials aiming to improve the treatment of cryptococcal meningitis, but the relationship of isolates from these countries to the global population is not yet described. Here, we address this knowledge gap by using Multilocus Sequence Typing to study the population of *Cryptococcus neoformans* var. *grubii* (CNVG) in Laos and Vietnam, with the specific aim of incorporating these populations into the wider global context. We found that, in most countries, a single lineage (family) of strains was responsible for most disease. The Vietnamese CNVG population was unusual in that 2 main lineages circulated at the same time. The Vietnamese CNVG population occupies a middle ground between Thailand/Laos in the west and China in the east. The differences in population structure moving from West to East are probably due to ecological differences. Disease in HIV uninfected patients was almost always due to members of a single family of strains (ST5).

## Introduction

*Cryptococcus neoformans* is the main etiological agent of cryptococcosis. It is of major significance in HIV patients and other individuals with immunosuppression, accounting for an estimated 223,100 incident cases of meningitis per year globally [1]. *C. neoformans* is an environmental saprophyte associated with bird guano and trees [2–5]. There is no human to human spread but human infection is believed to result from the inhalation of desiccated yeasts or spores[6]. While *C. neoformans* is commonly associated with immunosuppression, the related species *C. gattii* is associated with infection in immunocompetent hosts, especially in tropical and subtropical regions [7–11]. An outbreak of *Cryptococcus gattii* disease in mostly immunocompetent patients has been reported from Vancouver Island, British Columbia, Canada since 1999 [12]. Subsequent cases reported from Washington and Oregon suggest that *C. gattii* is also established in in the Pacific Northwest of America [13]. In Asia, the majority of cases of cryptococcal meningitis are due to *C. neoformans* var. *grubii* (CNVG)[14–17]. Here, in addition to the strong association with HIV infection, CNVG is also reported to affect apparently healthy individuals[7,18]. Such cases have been reported from China, Japan, Korea and Vietnam, and are due to a clonal group of CNVG [14,19–22].

Southeast Asia is a geographically and ecologically diverse area known for its high species density[23,24]. The south of the region has a hot tropical climate with dry and wet seasons; the north is temperate with hot summers and cold winters. The Annamite mountain range, covered in dense rain forests, rises to over 3000 meters and runs the length of Vietnam, forming a barrier separating the Western regions of Thailand, Laos and Cambodia from Vietnam and southern China. Despite the diverse environmental niches, the vast majority of clinical and environmental CNVG isolates from South, East and Southeast Asia, have been found to be serotype A, molecular type VNI, mating type MATα, suggesting a clonal population structure [25]. The dominance of MATα isolates from both the environment and human patients also suggests a clonal mode of expansion. Previous studies suggest there has been long distance dispersal of the species out of Africa [5,26]. Multilocus Sequence Typing (MLST) epidemiological studies from East Asia have shown limited genetic diversity among the majority of environmental and clinical isolates [19,27,28]. Despite the evidence of global dispersion at the species level[29,30], different geographical areas have distinct population structures, particularly in relation to the predominant sequence types[10]. Previous studies suggested that though in relatively close geographical proximity, the population structure of CNVG from Thailand and China are significantly different[31,32]. However, few data exist from other countries in the Southeast Asia region.

Here, we describe the population structure *of C. neoformans* var. *grubii* from Laos and Vietnam, two neighboring countries separated by the Annamite range, and integrate these data into the wider global context by including published data. Our specific goals were (i) to describe the genetic structure of the Vietnamese and Laotian CNVG population using MLST, (ii) to set the population structure within the global context, and (iii) to investigate links between infecting CNVG genotypes and host HIV-status.

## Methods

### Ethics statement

All studies from which samples were derived were approved by the Hospital for Tropical Diseases Ethical Review Board or the National Ethics Committee for Health Research, Ministry of Health, Lao PDR, and either the Oxford Tropical Ethics Committee, University of Oxford, or the Ethics Committee of the Liverpool School of Tropical Medicine. All patients, or their responsible next of kin, gave written informed consent to enter clinical studies.

### Patients and isolates

The Vietnamese isolates (total n=136) included in this study were clinical isolates from the cerebrospinal fluid (CSF) of patients enrolled in a randomized controlled trial of antifungal therapy in HIV-infected patients (n=97) between 2004 and 2011, and a prospective, descriptive study of HIV-uninfected patients with central nervous system (CNS) infections (n=39) enrolled between 1998 and 2009 [14,25]. The Laotian isolates were from 81 patients with cryptococcal meningitis (67 HIV-infected and 14 HIV-uninfected) consecutively admitted to Mahosot Hospital, Vientiane, between 2003 and 2015.

### DNA extraction and Restriction Fragment Length (RFLP)

Total genomic DNA was extracted from all organisms using the Masterpure Yeast DNA kit (Epicentre Biotechnologies, Madison, WI, USA) according to the manufacturer’s instructions. Restriction fragment length polymorphism analysis (RFLP) of the orotidine monophosphate pyrophosphorylase (URA5) gene was used to confirm species/varietal status of all *C. neoformans* var. *grubii* isolates [33]. PCR of the URA5 gene was conducted in a final volume of 50 μL. Each reaction contained 50 ng of DNA, 1X HotStart PCR buffer (NEB, USA), 0.2 mM each of dATP, dCTP, dGTP, and dTTP (Roche Diagnostics GmbH), 2 mM magnesium acetate, 0.5 U HotStart Taq DNA polymerase (NEB, USA), and 0.15 μΜ of each primer URA5 (5'-ATG TCC TCC CAA GCC CTC GAC TCC G - 3') and S01 (5'-TTA AGA CCT CTG AAC ACC GTA CTC - 3'). PCR was performed in an Nexus SX1 thermal cycler (Eppendorf, USA) at 95°C for 15-min, followed by 35 cycles of 20s denaturation at 94°C, 40s annealing at 61°C, and 1 min extension at 72°C, followed by a final extension cycle for 3 min at 72°C. 10 μL of PCR products were double digested with Sau96I (0.2 U/μL.) and Hhal (0.8 U/μL.) for 3 h at 37°C, followed by a final 10min incubation at 60°C. Restriction products are separated by 3% agarose gel electrophoresis at 100 V for 5 h. RFLP patterns were assigned visually by comparing them with the patterns obtained from the control strains (VNI-VNIV and VGI-VGIV, See Supplementary table S2) kindly provided by Prof. Wieland Meyer, University of Sydney (for details see supplementary materials).

### Multilocus Sequence Typing (MLST)

Seven MLST loci (CAP59, GPD1, IGS1, LAC1, PLB1, SOD1, and URA5) were amplified and sequenced following the procedures of the International Society for Human and Animal Mycology (ISHAM) MLST consensus typing scheme for *C. neoformans* (http://mlst.mycologylab.org) [34]. Sequencing was performed using BigDye v3.1 Chemistry (Applied Biosystems, CA, USA) on an ABI 3130 Genetic Analyzer (Applied Biosystems, CA, USA). Both forward and reverse amplicons of each locus were sequenced. Consensus sequences were manually edited using ContigExpress and aligned in AlignX, implemented in VectorNTI Suite 7.0 [35]. A single Allele Type (AT) number was assigned to each of the seven loci by comparing consensus DNA sequences with the ISHAM database, resulting in a seven-number allelic profile for each isolate. The allelic profiles defined the corresponding STs.

MLST profiles and DNA sequences at each MLST loci for isolates from regions other than Laos and Vietnam were obtained from NCBI. For the global analysis, we used data reported by Simwami *et al*. [31], Mihara *et al*. [20], Cogliati *et al*. [36], Khayhan *et al*. [10], Chen *et al*. [37] and Dou et al. [38].

### Phylogenetic analyses

For global comparison, information on MLST genotypes, patient HIV status, and source of isolation for 617 isolates from other countries across Asia and Southern Africa were obtained from NCBI as previously described[10,20,31,36,37].These consisted of clinical and environmental *C. neoformans* var. *grubii* isolates from Southeast Asia (Indonesia, N=41 and Thailand, N=222); East Asia (Japan, N=38 and China, N=132); South Asia (India, N=61), Middle East (Qatar, N = 5 and Kuwait, N=10) and Africa (Botswana, N=141).

First, we determined the best DNA substitution model for the concatenated dataset using MEGA v6.0.6 [39]. The Kimura 2-parameters model with gamma distribution was selected as the best fit model using the Bayesian information criteria (BIC) incorporated in MEGA v6.0.6[39,40]. To evaluate patterns of evolutionary descent among genotypes according to their source and geographic region, the allelic profile of the Vietnamese, Laos and global dataset were applied to the goeBURST algorithm in PHILOVIZ 2.0 software available at http://www.phyloviz.net[41,42]. A group founder was defined as the sequence type with the most number of linked single locus variants (SLV). The concept of a clonal complex (CC) was adopted when a SLV linkage with the founder ST was observed.

### MDS clustering

Genetic distance (Fst) and geographical distance were visualized by the non-metric multidimensional scaling method (MDS). Fst were calculated from concatenated sequences of the 7 MLST housekeeping genes using DnaSP v5 [43]. MDS calculation was performed using RStudio Version 0.98.1103 built with R-3.4.0 (https://www.rstudio.com and https://www.r-project.org). Network analysis and visualization was conducted using the R package igraph (version 1.1.2).

## Results

### MLST reveals different sequence type distribution pattern between Laos and Vietnam

All isolates from Vietnam were previously determined as molecular type VNI by RFLP, mating type α [25]. The 136 Vietnamese isolates consisted of 13 different sequence types (ST) of which ST5 (n=66; 48%) and ST4 (n=32; 23%) were the most prevalent, followed by ST6 (n=12; 9%), ST93 (n=8; 6%), ST32 (n=7; 5%), ST39 (n=3; 2%) and 7 less common STs with less than 3 isolates each (see Table 1). Of 13 STs detected in Vietnam, were previously undefined in the MLST database: ST306, ST338, ST339 and ST340. These 4 STs were singletons, each accounting for <1% of the total number of isolates. All 81 Laotian isolates also belonged to the RFLP-defined VNI group. Among them, ST4 and ST6 accounted for over 40% each of the total number of isolates, and were 4-fold more prevalent than ST5; (ST4, n=35 (43%); ST6, n=32 (40%); ST5, n=9 (11%), Fig 1A, Table 1). Less than 3% of isolates from Laos (n=2) were ST93. Of note, ST306 was found in both Vietnam and Laos. ST32 was not identified among the Laos isolates.

**Table 1.**
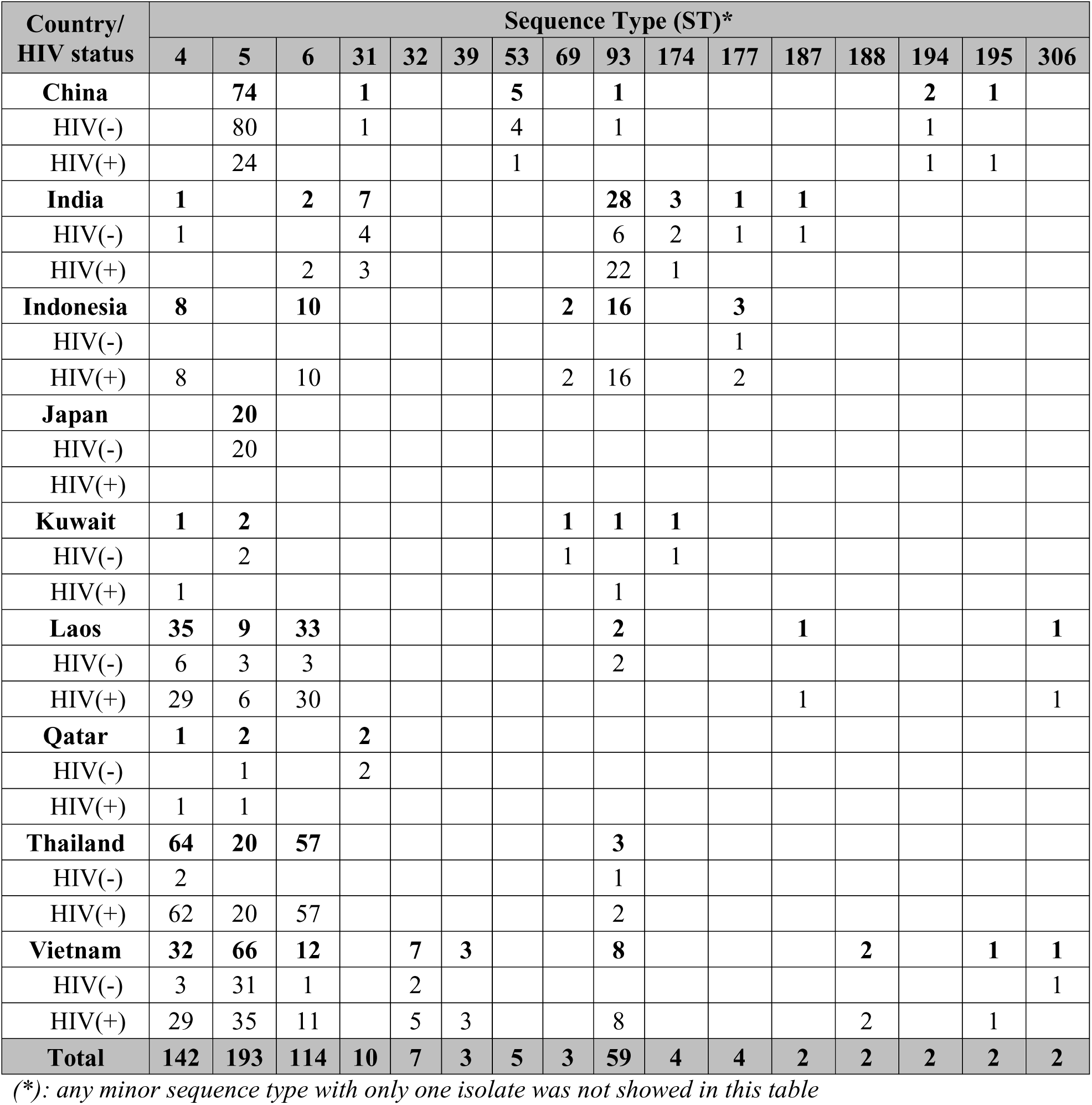
Distribution of major Sequence Type (ST) in the Asia/Middle East region according to country of origin and HIV-status.

**Fig 1.**
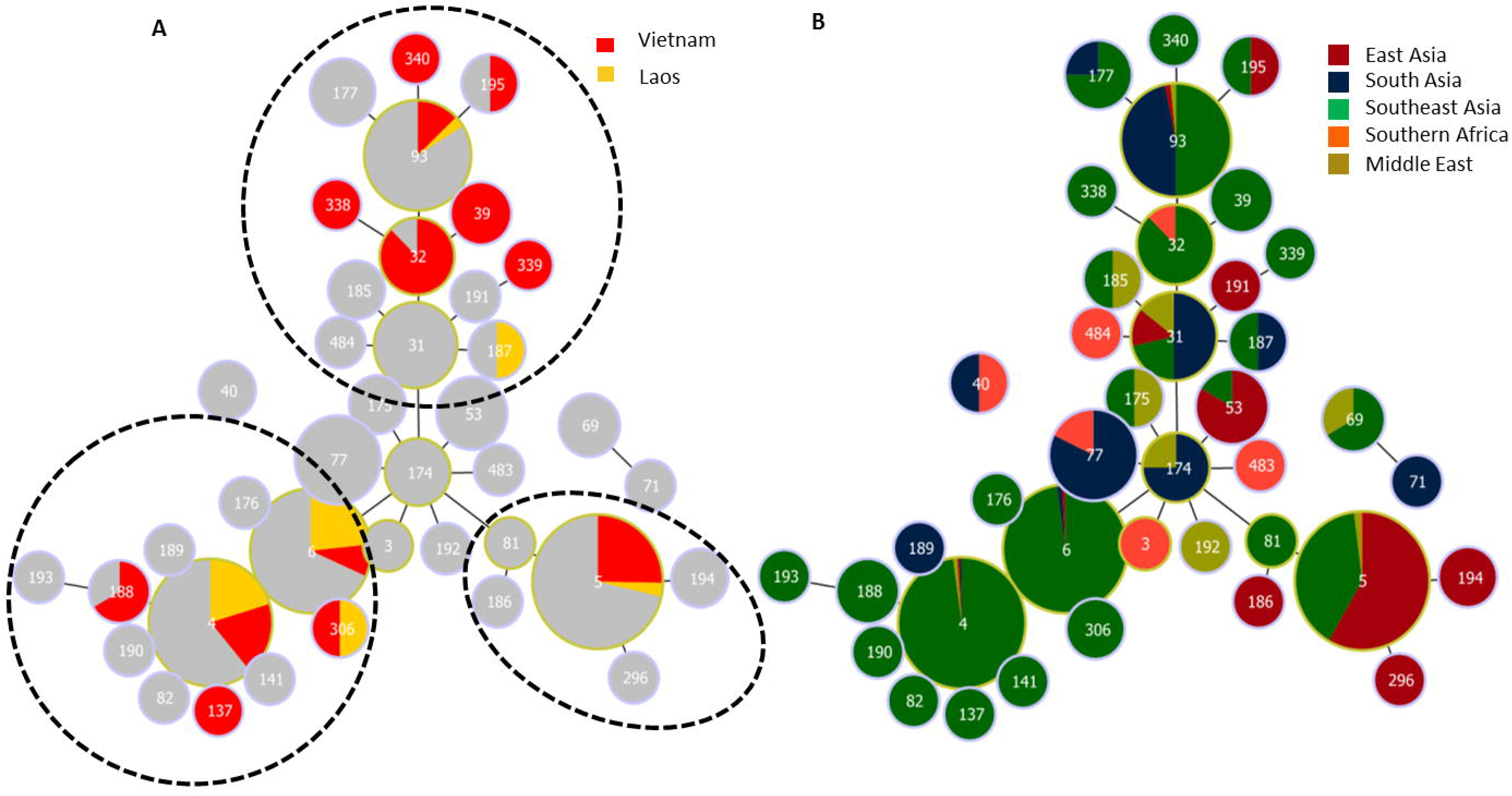
goeBurst analysis integrating the Laotian and Vietnamese *C. neoformans* var. *grubii* isolates into the major Asian clonal complex. Minimum spanning tree showing the major Asian clonal complex. **(A)**: Laos and Vietnam as part of the global population. 3 main subgroups of the major Asian clonal complex are indicated by dotted circles. **(B)**: the Asian clonal complex broken down into geographical regions: East Asia (China and Japan), Southeast Asia (Vietnam, Laos, Thailand and Indonesia), South Asia (India), plus the reference groups Middle East (Qatar and Kuwait) and Southern Africa (Botswana). A group founder was defined as the sequence type with the most number of linked SLVs. The concept of a clonal complex (CC) was adopted when a SLV linkage with the founder ST was observed. All Asian/Middle Eastern STs were interlinked by only SLVs and formed a major clonal complex around the group founder, ST174. ST174 were only found in South Asia and the Middle East. 3 sub-groups deriving from ST174 were specified, each of which was associated with a geographical region.

### Correlation between infection with ST5 and HIV status

Of the 39 HIV-uninfected patients with cryptococcosis in Vietnam, 31 infections were due to ST5 (79%); the remaining 8 patients were infected with ST4, ST32, ST6 and ST306. Three of 39 HIV-uninfected patients had other underlying disease; all these patients had non-ST5 infections. 36% of cases in HIV-infected patients (N=35) were due to ST5 while ST4, ST6 and other non-ST5 sequence types accounted for the remaining 64% (N=63). Compared to all other STs, ST5 in Vietnam was significantly associated with HIV-uninfected patients (Odds Ratio 6.87 [95%CI: 2.72 to 19.26], *p*<0.0001, Fisher’s exact test). In contrast, 3 out of 14 (21%) cases in HIV-uninfected patients in Laos were due to ST5; in HIV infected patients ST5 accounted for 6 of 67 (9%) cases of meningitis (Odds Ratio 2.72 [95%CI: 0.38 to 15.20], *p*=0.18, Fisher’s exact test).

We placed the association of HIV serostatus in the Vietnamese and Laotian dataset in the global context by gathering previously described data from Asia and Africa [10,37]. We excluded the 47 clinical isolates in the global dataset with unknown HIV status. Of 198 HIV uninfected patients, 137 (69%) were infected with ST5 isolates; of 477 HIV infected patients, 87 were infected with ST5 isolates (Odds Ratio 10.02 [95%CI: 6.76 to 15.01], *p*<0.0001, Fisher’s exact test), East and Southeast Asia accounted for the majority of clinical and environmental ST5 isolates worldwide (n=256; 98%). The relative proportion of ST5 isolates from each country decreased along an arc from Northeast to Southwest: Japan (n=36; 95%), China (n=116; 87.9%), followed by Vietnam (n=66; 48.5%), Thailand (n=29; 13.1%) and Laos (n=9; 11.1%). ST5 accounted for 20% (n=2) and 40% (n=2) of the total number of isolates from Kuwait and Qatar, respectively. Only one single ST5 isolate was reported in the Botswana dataset.

### goeBurst analysis and geographical distribution

To further elucidate how the Vietnamese/Laotian populations fit into the wider global context, we expanded the analysis to include previously published data from different Asian regions. goeBurst analysis placed all Asian and Middle Eastern *C. neoformans* var. *grubii* isolates (725/866; 84%) in a clonal complex (Fig 1B), with VNI Vietnamese/Laotian populations present across the diversity of the complex (Fig 1A). The Asian clonal complex was separated from the majority of Botswana isolates at all 7 loci. ST174 was identified as the major group founder of the Asian clonal complex when analyzed by both parsimony (Fig 1) and maximum likelihood analyses (Fig S2). The main Asian clonal complex included 3 main goeBurst sub-groups arising from ST174; sub-group 4 (ST4/ST6), sub-group 5 (ST5) and sub-group 93 (ST31/93) (Fig 1A). No sub-groups were assigned to the Botswana isolates (N=141) which were mostly distant from the Asian clonal complex.

We observed an association between geographical location and STs (Fig 1B). Most isolates from South Asia (India) and the Middle East were ST174 and variants from sub-group 93. Sub-group 4 was highly prevalent in Southeast Asia; 318 out of 326 (98%) sub-group 4 isolates were from Southeast Asia. 256 out of 261 (98%) isolates from sub-group 5 were from East or Southeast Asia. Sequence type 93 was the most widely distributed, being reported from Southeast Asia (n=31; 6% of isolates from this region, like all percentages reported in this sentence), South Asia (n=29; 48%), East Asia (n=1, 0.6%) and the Middle East (n=1; 7%) (See Fig 1B).

### Genetic distance between *C. neoformans* population from different geographical regions

We employed the fixation index (*Fst*) to evaluate pairwise genetic differentiation between different geographical regions. Multidimensional scaling (MDS) was carried out to provide a 2-D visualization of the *Fst* distance matrix (Table 2 and Fig 2). Botswana, being structurally diverse, is in stark contrast to any other countries and therefore was removed from the analysis. On the other hand, due to the apparent skewness of ST5 toward HIV-uninfected, apparently immunocompetent patients, the *Fst* analysis only included isolates from HIV-infected patients from Laos, Thailand, Vietnam, Indonesia and India. Very few isolates from Japan, Kuwait or Qatar were known to be from HIV-infected patients and thus isolates from these countries were also excluded from this analysis. The MDS plot revealed that while the Indian CNVG population was genetically distant to those in Southeast Asia, this population was most closely related to the Indonesian CNVG population (*Fst*=0.235, compared with *Fst* ranging from 0.535 to 0.777 for Vietnam, Thailand and Laos). The Indonesian CNVG population was most closely related to the CNVG populations in Vietnam (*Fst* = 0.153) and India (*Fst*=0.235) while CNVG populations from Laos and Thailand were genetically closer to each other (*Fst* = 0) than any other countries in the region. The CNVG population in Vietnam, while being placed closer to Indonesia in the MDS analysis, appeared as intermediate between China in East Asia (*Fst*=0.207) and Laos/Thailand in Southeast Asia (*Fst*=0.178 and *Fst*=0.151, respectively). In fact, the CNVG population in China is more closely related to the CNVG population in Vietnam (*Fst*=0.207) than either Laos or Thailand (*Fst*=0.583 and *Fst*=0.534, respectively).

**Table 2.** Genetic distance matrix (Fst) of Southeast Asian countries, with reference to
Botswana.

|  | Vietnam | Laos | Thailand | China | India | Indonesia |
| --- | --- | --- | --- | --- | --- | --- |
| Vietnam | 0 | 0.178 | 0.151 | 0.207 | 0.535 | 0.153 |
| Laos |  | 0 | 0.000 | 0.583 | 0.777 | 0.381 |
| Thailand |  |  | 0 | 0.534 | 0.767 | 0.373 |
| China |  |  |  | 0 | 0.747 | 0.450 |
| India |  |  |  |  | 0 | 0.235 |
| Indonesia |  |  |  |  |  | 0 |

**Fig 2.**
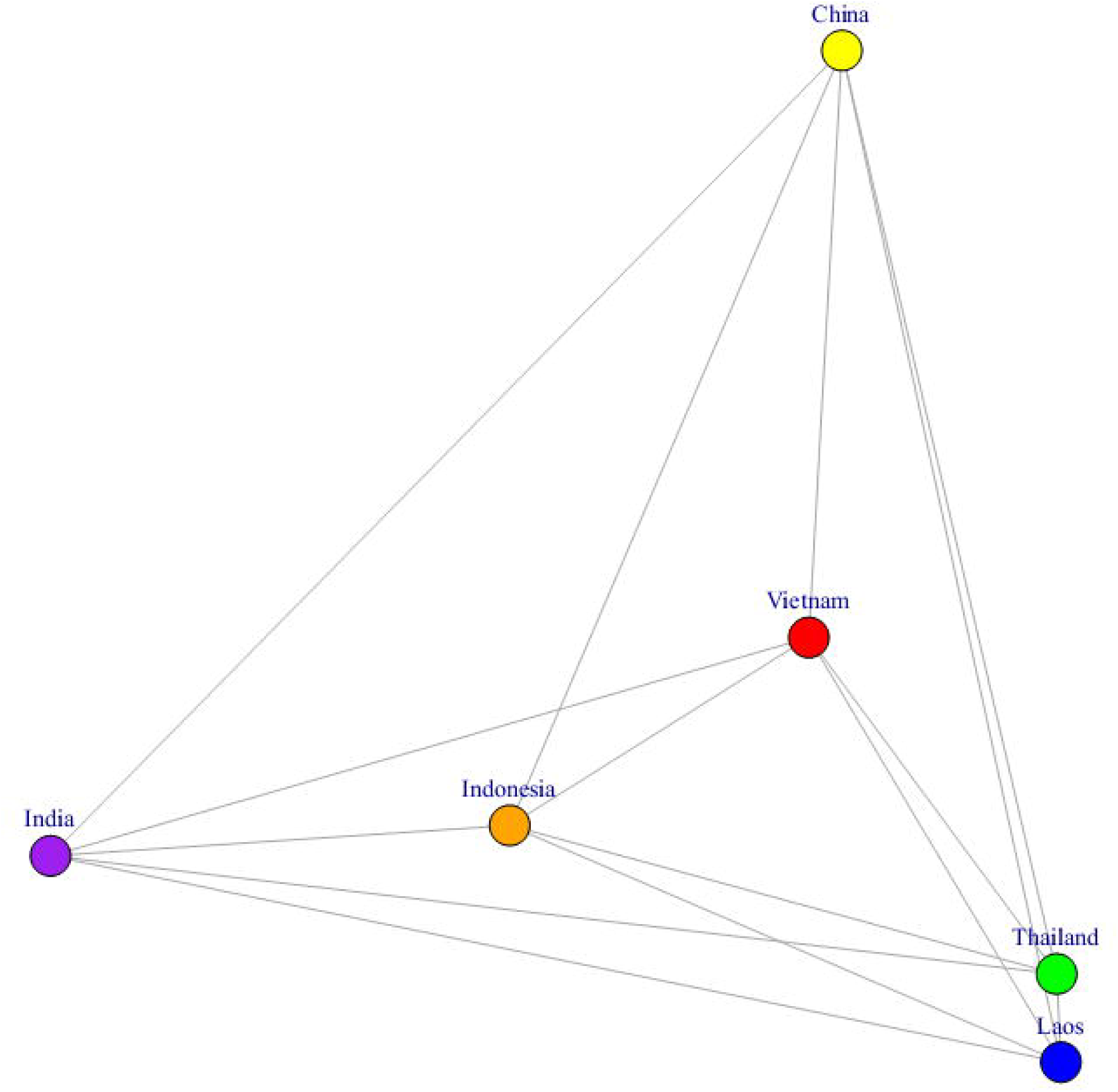
Mutidimensional scaling plot of genetic distances (Fst) of global *C. neoformans* isolates from HIV-infected patients (excluding the Botswana, Japan, Kuwait and Qatar). Vietnam appeared to represent an intermediate between Laos/Thailand and Indonesia. Indonesia is the only country in Southeast Asia to share the most similarity with India from South Asia, possibly due to the uniquely high proportion of ST93.

## Discussion

The population structure of *C. neoformans* var. *grubii* has been described from a number of Asian countries [10], including Thailand[17,31], India[44], China[45–8], Japan[20] and Korea[19]. However, data from Vietnam and Laos have been lacking. To address this, we analyzed 217 isolates from the 2 countries (136 isolates from Vietnam and 81 isolates from Laos) with MLST and used publicly available MLST data to set these populations in the global context. A strength of our dataset is that it contains significant numbers of strains from both HIV infected and uninfected patients. In contrast, reports from China and East Asian countries have focused on clinical isolates from HIV-uninfected patients, while studies from Thailand and other Southeast Asian countries including Singapore[49], Malaysia[50] and Indonesian[51], present isolates predominantly from HIV-infected patients. These differences between the immune statuses of the sources of clinical isolates complicate interpretation of the data, but also mean that Vietnam is in a unique position to compare CNVG population structure between these two patient groups.

In China, ST5 is the dominant ST, accounting for over 90% of infections [15,27,38]. While it is prevalent in both immunocompetent and immunocompromised patients, relatively few cases of HIV-associated disease have been reported [38]. It is not clear why there are so few cases of HIV-associated disease. Despite the relatively low prevalence of HIV in China (estimated at <0.1% of the adult population in 2015 [52]), given the numbers of cases of non-HIV-associated disease reported, this may represent bias in study design, HIV testing or reporting. In contrast to China, the CNVG population from Thailand is more diverse according to MLST, with 14 STs reported, among which ST4 and ST6 are most common, outnumbering ST5 infections 6 to 1 [10,17,31]. The Thai data are notable for the lack of CM cases reported from HIV-uninfected patients. This could represent under-reporting of non-HIV associated disease, or be due to a low prevalence of ST5 isolates in the environment. As we have shown here and previously, the ST5 lineage is associated with apparently immunocompetent patients in Vietnam [25,53]. ST93 was the most prevalent sequence type in India, Indonesia, Uganda and Brazil (varying between 48% and 77%) [10,54,55], but accounted for less than 4% of isolates from Vietnam and Laos.

We found that the population structure of isolates from Laos appears similar to that in Thailand, with ST4 and ST6 (both belonging to sub-group 4) accounting for the majority of disease. The dominance of one particular sub-group in Thailand/Laos and China contrasts with Vietnam, where ST5, ST6 and ST4 (belonging to two different sub-groups) are co-dominant. It is possible that the prevalence of ST5 in Thailand and Laos is under-reported due to lack of isolates from HIV uninfected patients; thus an association between ST5 and HIV-uninfected patients could not be confirmed in these two countries. However, the relatively high prevalence of ST5 strains in the Vietnamese dataset is not solely driven by isolates from HIV uninfected patients - this ST still represents >35% of cases of disease in HIV infected patients. Thus it appears that Vietnam represents an intermediate zone between Laos/Thailand to the west, and China to the east, consistent with its geographical location.

The differences in population structure we see moving from west to east are likely driven by ecological and/or human factors. Ecological differences between the regions could select for enrichment of particular STs if there are differences in the relative fitness of the STs in the varying ecological niches. Potential differences across the region include shared differences in climate (tropical savannah through mountainous rain forest to temperate), and differences in elevation (for example, Chiang Mai in Northern Thailand and Vientiane in Northwestern Laos are at significantly higher elevation than Ho Chi Minh City and much of Southeastern China (170-300 meters versus 10-20 meters above sea level). Other than elevation and temperature, localized differences in soil biogeochemistry, environmental predators [56] such as amoebae or tree species may also be relevant. For example, Laos, Cambodia, Vietnam and China are among Asian countries where there is significant contamination of groundwater with high, potentially harmful levels of arsenite (>10 μg/L, WHO guidelines)[57,58]. This distribution to some extent follows that of the ST5 lineage. Recently it has been shown that ST5 *C. neoformans* var. *grubii* isolates possess tandem repeats of the arsenite efflux transporter (Arr3) [59]. The greater the number of these repeat elements the higher the resistance to toxic arsenic containing compounds [59]. Correlating Arr3 copy numbers in environmentally sourced strains with environmental arsenic concentrations would address this hypothesis, although isolating CNVG from the environment is technically challenging.

In addition to ecological factors, the varying prevalence of lineages in particular human populations could be a result of differential host susceptibility to those lineages. Such human genetic factors have previously been associated with susceptibility to tuberculosis [60]. Regarding the host population structure in Southeast and East Asia, the dominant ethnic group in Vietnam (Kinh, >80%), is genetically closer to ethnic groups in Northern Thailand than to the Han Chinese from Beijing or Southern China [61] (also see Fig S1). Thus, while the Vietnamese human population is genetically closer to that from Northern Thailand than that from China, we demonstrated the contrary pattern regarding the populations of clinical isolates of CNVG, suggesting that the diversity of the pathogen population is not driven by host susceptibility. Nevertheless, it remains possible that subtle human genetic polymorphisms exist that result in differential susceptibility to different cryptococcal species/lineages.

Our observation suggest that the populations of *C. neoformans* var. *grubii* found in many geographical regions across Asia and the Middle East arose from the clonal expansion of a single genotype, ST174, and is consistent with a previous report [55]. This ST174 clonal expansion model is supported by goeBurst analysis which shows that ST174 is the most parsimonious candidate as the group founder to subgroup 4, 5 and 93, which account for over 90% of the Southeast Asian isolates. Maximum likelihood phylogeny also supports this finding. ST174 was not observed in either Vietnam or Laos but was present in previously published datasets from the Middle East, with 3 isolates from India and 1 isolate from Kuwait.

## Conclusion

Our study addressed the lack of knowledge of the molecular epidemiology of *C. neoformans* var. *grubii* from Vietnam and Laos in Southeast Asia. We found a change in predominant STs as one moves from western to eastern longitudes, that the Vietnamese *C. neoformans* var. *grubii* population appears intermediate between Thai and Chinese populations, and is unique in there being co-dominant subgroups. Our data suggest that ST174 is the putative most recent common ancestor for the majority of CNVG sequence types in Asia. The *C. neoformans* var. *grubii* population in Asia present in this study appeared mostly clonal, which is similar to previous reports. Most studies focusing on VNI, including ours, are weakened in that while cryptococcosis results from the inhalation of infectious propagules from the environment, few environmentally sourced isolates have been typed, and thus the true diversity of the species in any country may be underestimated. More efficient environmental isolation techniques and systematic sampling would allow us to understand the true species diversity in the region and could improve understanding of the basis of cryptococcal virulence as well as bottleneck events leading to the rise of pathogenic lineages.

## Acknowledgements

We thank Dr. Lu Dongsheng from Dr. Shuhua Xu’s group at the Max-Planck Independent Research Group on Population Genomics for providing human WGS SNP data from across Asia.

## Supporting information

**Figure S1.**
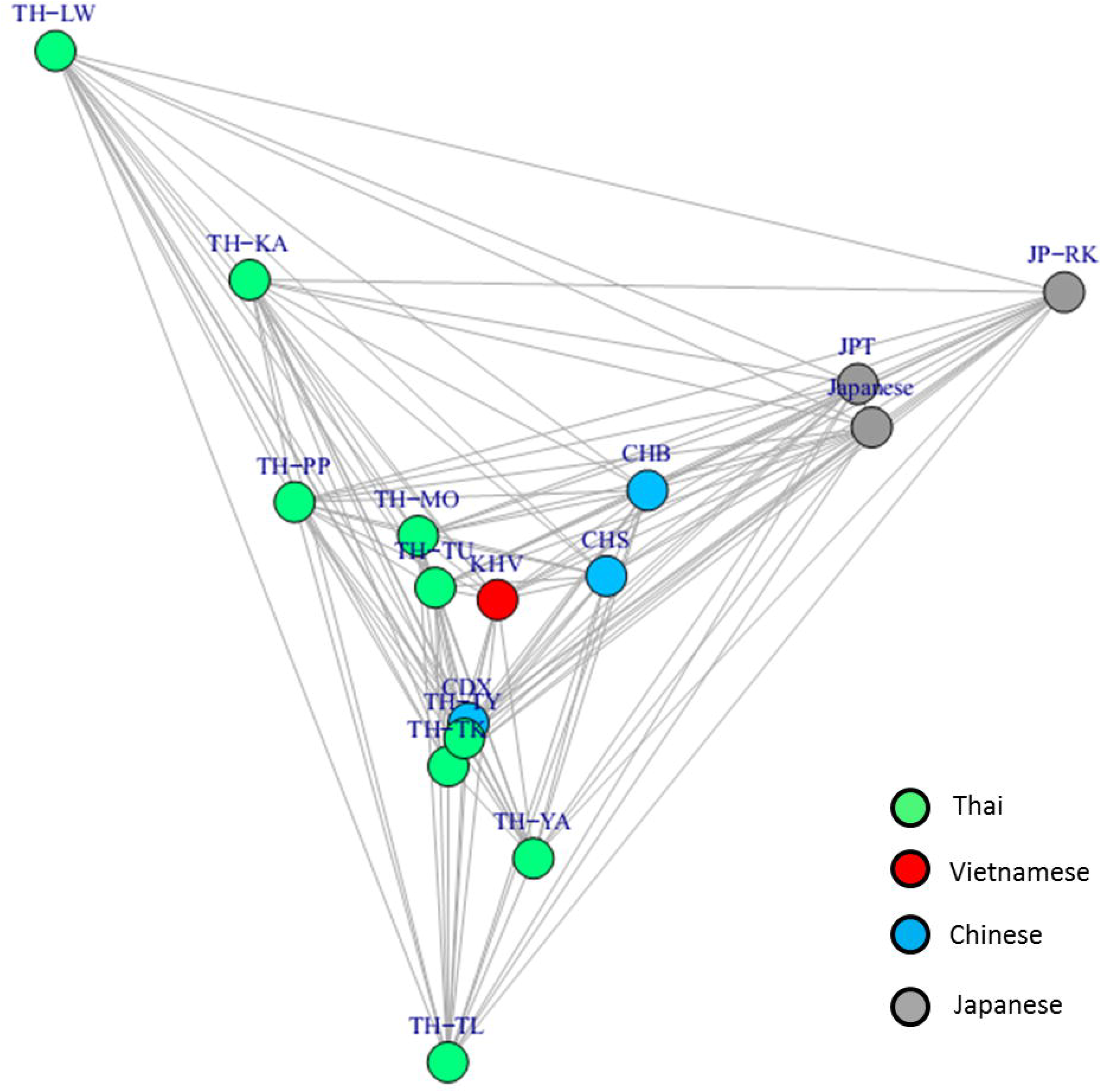
Genetic distance between human populations from Japan, China, Vietnam and Thailand. MDS plot showing SNPs data from WGS of different Asian populations from Vietnam, Japan, China and Thailand. Thai groups include ΤΗ-PP (Plang), ΤΗ-PL (Palong), TH-TK (Tai Kern), TH-TL (Tai Lue), ΤΗ-MO (Mon), TH-LW (Lawa), TH-KA (Karen), TH-YA (Yao), TH-TN (H’Tin), TH-MA (Mlabri), ΤΗ-ΤΎ (Tai Yong), TH-TU (Tai Yuan), TH-HM (Thai H’mong). Japanese groups include: JPT (Japanese from Tokyo), JP-RK (Ryukyuan) and Japanese (data from Human Genome Diversity Panel-CEPH). Chinese groups include Han Chinese from Beijing (CHB) and Southern China (CHS). Dai Chinese from Xishuangbanna was previously classied as a distant Southeast Asian population. Kinh Vietnamese from 2 most populous cities in Vietnam, Ho Chi Minh and Hanoi, were designated as KHV. Lu Dongsheng (Max Planck Independent Research Group on Population Genomics) compiled and provided human genome SNP data from the Human Genome Diversity Panel-CEPH, the HUGO Pan-Asian SNP Consortium and the 1000 Human Genome Project.

**Figure S2.**
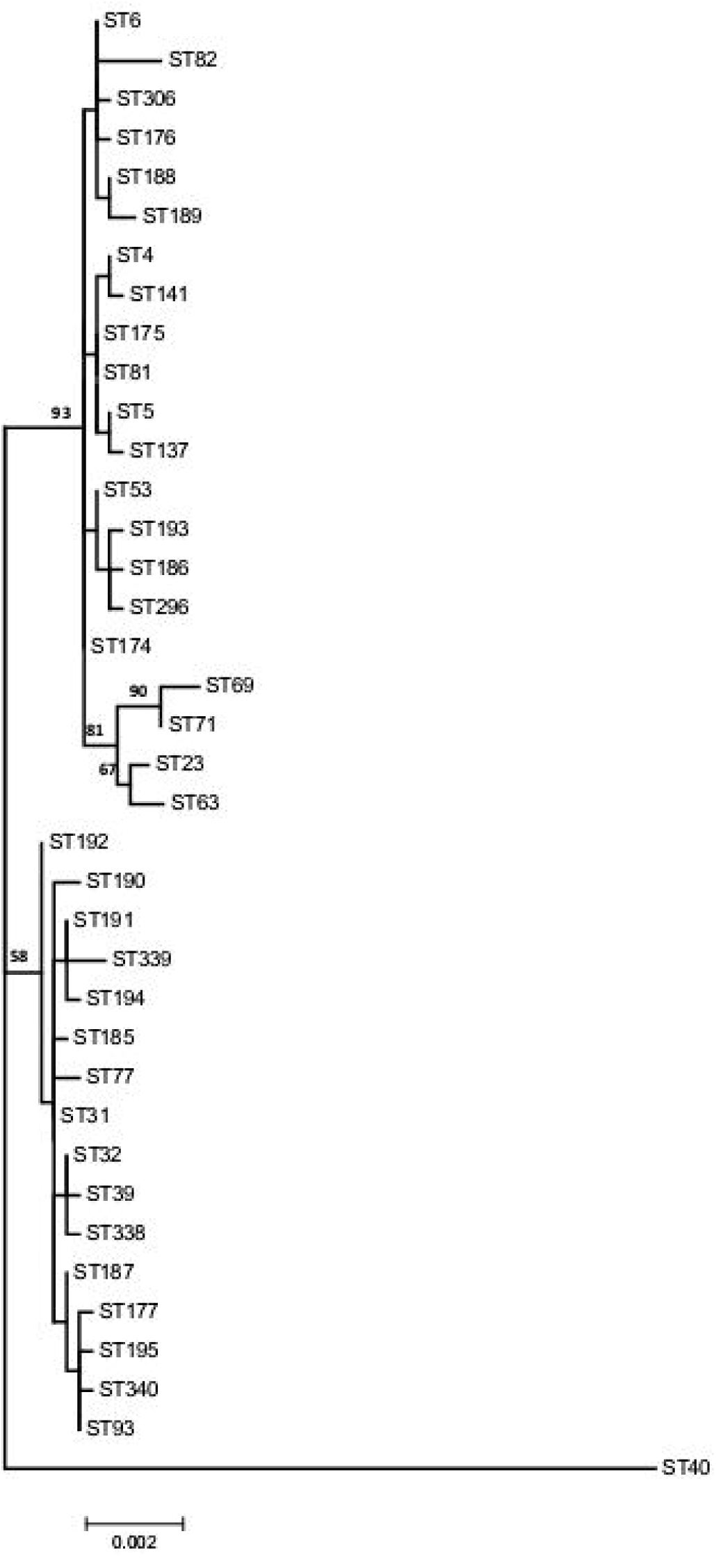
Molecular Phylogenetic analysis by Maximum Likelihood method. The evolutionary history was inferred by using the Maximum Likelihood method based on the Kimura 2-parameter model. The tree with the highest log likelihood (-6364.7916) is shown. The percentage of trees in which the associated taxa clustered together is shown above the branches. Initial tree(s) for the heuristic search were obtained automatically by applying Neighbor-Join and BioNJ algorithms to a matrix of pairwise distances estimated using the Maximum Composite Likelihood (MCL) approach, and then selecting the topology with superior log likelihood value. A discrete Gamma distribution was used to model evolutionary rate differences among sites (5 categories (+*G*, parameter = 0.1000)). The rate variation model allowed for some sites to be evolutionarily invariable ([+/], 48.9615% sites). The tree is drawn to scale, with branch lengths measured in the number of substitutions per site. The analysis involved 38 nucleotide sequences. There were a total of 3996 positions in the final dataset. Evolutionary analyses were conducted in MEGA6. Bootstrap values over 50% were shown.

**S1 Table. goeBURST subgroups**

**S2 Table. Isolates info**

**S3 Table. MLST Allelic profiles**

**S4 Table. goeBURST subgroups and HIV-status according to countries**

**S5 Table. ST and HIV status according to countries**

